# Thermal stress-based pathway engineering and bioprocess optimization of *Corynebacterium glutamicum* for *de novo* hydroxyectoine production

**DOI:** 10.64898/2026.09.17.752347

**Authors:** Luciana Fernandes Brito, Nathalie van Assel, Fernando Pérez-García

**Affiliations:** Department of Biotechnology and Food Science, Faculty of Natural Sciences, NTNU, Trondheim, Norway

**Keywords:** Hydroxyectoine, ectoine, *Corynebacterium glutamicum*, thermal stress, synthetic operon, bioprocesses development

## Abstract

**Background:** Hydroxyectoine is a high-value compatible solute with applications in cosmetics, healthcare, and biotechnology. Its microbial production has traditionally relied on halophilic organisms requiring high-salinity cultivation, motivating the development of non-halophilic production hosts. Corynebacterium glutamicum is an attractive platform because of its industrial robustness and high metabolic capacity through the aspartate-family amino acid pathway.

**Results:** A synthetic hydroxyectoine pathway was constructed by combinatorially varying the predicted translation initiation rates of four translational units comprising *ask*, *ectAB*, *ectC,* and *ectD*. A library of *C. glutamicum* transformants was screened at 40 °C using hydroxyectoine-associated thermoprotection as a functional selection principle. Fast-growing variants generally displayed increased hydroxyectoine formation and higher hydroxyectoine-to-ectoine ratios than a strain carrying the native pathway configuration. The selected combination reached a hydroxyectoine-to-ectoine ratio of 4.7 and was transferred into the lysine-producing strain DM1729SL in order to increase precursor availability and hydroxyectoine production. Carbon-limited fed-batch cultivations showed that both temperature and dissolved oxygen influenced production. The best performance was obtained at 30 °C and 50% relative dissolved oxygen, yielding 12.9 g/L hydroxyectoine with a yield of 0.161 g/g glucose and a volumetric productivity of 0.26 g/L/h after glucose depletion. A subsequent hypoosmotic downshock increased extracellular hydroxyectoine recovery to 14.5 g/L.

**Conclusions:** Thermal stress-based combinatorial pathway balancing enabled efficient *de novo* hydroxyectoine production in *C. glutamicum*. Integration of precursor-enhanced host metabolism, controlled oxygen supply, fed-batch cultivation, and osmotic downshock resulted in a substantial improvement over previously reported *de novo* hydroxyectoine production in this host and establishes *C. glutamicum* as a promising platform for further hydroxyectoine process development.

## 1. Introduction

Microorganisms exposed to osmotic, thermal, desiccation, and other environmental stresses frequently accumulate compatible solutes to preserve cellular function [1]. Among these compounds, ectoine and its hydroxylated derivative, 5-hydroxyectoine, hereafter referred to as hydroxyectoine, are cyclic non-proteinogenic amino acid derivatives produced by diverse bacteria and some archaea [1]. Both compounds stabilize proteins, membranes, and other macromolecular structures by promoting preferential hydration and maintaining the organization of the surrounding water network [2,3]. These protective properties have led to their application as high-value ingredients in cosmetics, healthcare products, and biotechnology [4]. Hydroxyectoine is particularly attractive because the additional hydroxyl group can provide enhanced protection against thermal and desiccation stress in comparison with ectoine [1,5].

The biosynthesis of hydroxyectoine originates from L-aspartate-β-semialdehyde, an intermediate of the aspartate-family amino acid pathway (Fig. 1). EctB converts L-aspartate-β-semialdehyde into L-2,4-diaminobutyrate using glutamate as cofactor. L-2,4-Diaminobutyrate is subsequently acetylated by EctA and cyclized by EctC to form ectoine. Hydroxyectoine is then generated through the stereospecific hydroxylation of ectoine by EctD [1]. This ectoine hydroxylase belongs to the non-heme Fe^2+^-and α-Ketoglutarate-dependent dioxygenase family and requires molecular oxygen, Fe^2+^, and α-Ketoglutarate for activity [6] (Fig. 1). Consequently, hydroxyectoine biosynthesis depends not only on expression of the heterologous pathway but also on the intracellular availability of precursor metabolites, oxygen, and cofactors (Fig. 1). Some organisms carry the *ectABCD* genes as a single operon like *Streptomyces coelicolor* [7], whereas others, such as *Pseudomonas stutzeri*, also include *ask* gene within the cluster [8]. The *ask* gene encodes aspartate kinase, which catalyzes the first committed step of the aspartate-family pathway using ATP as phosphoryl donor and supports hydroxyectoine biosynthesis by increasing metabolic flux toward the precursor L-aspartate-β-semialdehyde [9] (Fig. 1).

**Fig. 1.**
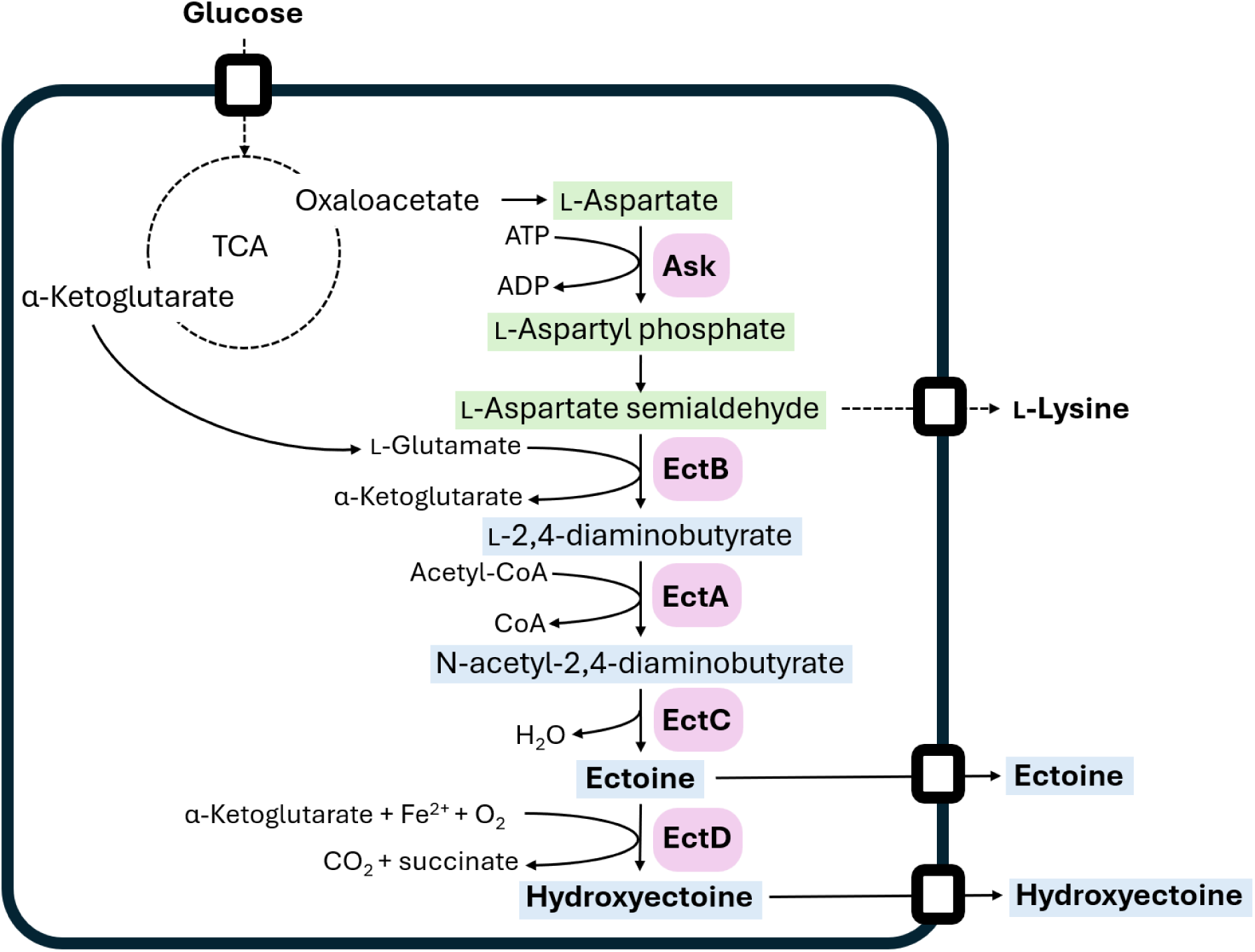
Hydroxyectoine biosynthetic pathway implemented and optimized in *C. glutamicum*. Green shadow metabolites: native metabolites involved in hydroxyectoine biosynthesis. Blue shadow metabolites: non-native metabolites involved in hydroxyectoine biosynthesis. Purple shadow enzymes: enzymes involved in the hydroxyectoine biosynthesis in *P. stutzeri* encoded in the cluster *ectABCD-ask*.

Traditional microbial processes for ectoine and hydroxyectoine production have commonly relied on natural halophilic producers like *Halomonas salina* or *Halomonas salifodinae* [10,11]. These organisms frequently require elevated salt concentrations to induce compatible-solute biosynthesis, which can increase corrosion, complicate downstream processing, and impose physiological limitations on high-cell-density cultivation [12]. Metabolic engineering of non-halophilic hosts provides an alternative approach that can decouple product formation from osmotic stress and enable production under more conventional fermentation conditions [12,13]. *Corynebacterium glutamicum* is an industrially established microbial production host with extensive applications in amino acid fermentation [14]. Its robustness, metabolic capacity, and naturally high flux through the aspartate-family pathway make it an attractive platform for producing ectoine-derived compounds [15,16]. Previous metabolic-engineering studies established heterologous ectoine and hydroxyectoine biosynthesis in *C. glutamicum* from L-aspartate-β-semialdehyde [13,16–19]. Efficient *de novo* hydroxyectoine production requires balanced expression of the ectoine biosynthetic enzymes and EctD. Excessive pathway activity may impose a metabolic burden and redirect carbon and precursor metabolites away from biomass formation, whereas insufficient EctD activity can result in ectoine accumulation and reduced hydroxyectoine selectivity [16,19]. Moreover, the expression level that is optimal for one enzyme may not be suitable for other pathway components. Combinatorial variation of translation initiation rates enables the exploration of this multidimensional expression space through compact pathway libraries, without requiring the separate rational construction of every possible expression configuration [20]. Linking pathway performance to a selectable or readily screenable cellular phenotype could further facilitate the identification of productive operon architectures. Hydroxyectoine is associated with protection against elevated temperatures, and disruption of *ectD* has been shown to compromise thermotolerance in the natural hydroxyectoine producer *Chromohalobacter salexigens* [21]. The established contribution of hydroxyectoine to thermoprotection suggested that intracellular hydroxyectoine formation could provide a functional basis for screening engineered strains under heat stress [21]. However, thermotolerance is a complex phenotype influenced by multiple physiological and experimental factors, including medium composition, cultivation conditions, metabolic burden, and the intracellular compatible-solute profile [22]. A thermal stress-based screening strategy therefore offers a practical approach for enriching promising pathway configurations, but subsequent direct quantification of ectoine and hydroxyectoine remains necessary to identify the most productive variants.

In this study, a synthetic hydroxyectoine pathway was developed in *C. glutamicum* by combinatorially varying the predicted translation initiation rates (pTIRs) of four translational units (TUs) comprising *ask*, *ectAB*, *ectC*, and *ectD*. The resulting plasmid library was screened at elevated temperature, exploiting hydroxyectoine-associated thermoprotection as a functional selection principle. Selected fast-and slow-growing *C. glutamicum* transformants were subsequently characterized with respect to operon architecture and extracellular ectoine and hydroxyectoine production, leading to the identification of plasmid pHect46. To enhance precursor availability, pHect46 was transferred into the lysine-producing strain DM1729SL, which carries increased metabolic capacity toward the aspartate-family pathway [17]. The effects of temperature, yeast extract supplementation, and host background on growth and product formation were then evaluated. Finally, hydroxyectoine production was investigated in carbon-limited fed-batch bioreactor cultivations at different temperatures and relative dissolved-oxygen setpoints. Together, this work establishes a thermal stress-based combinatorial engineering strategy and evaluates the metabolic and bioprocess factors governing *de novo* hydroxyectoine production by *C. glutamicum*.

## 2. Materials and Methods

### 2.1 Media and cultivation conditions

Table 1 details the specific bacterial strains and plasmids utilized throughout this study. Chemical reagents and general laboratory consumables were sourced from Sigma-Aldrich unless indicated otherwise. For cloning purposes, *Escherichia coli* DH5α served as the host organism, cultivated at 37 °C and 225 rpm in Lysogeny Broth (comprising 10 g/L tryptone, 5 g/L yeast extract, and 5 g/L NaCl) using either agar plates or liquid media. *C. glutamicum* strains were employed for heterologous expression. Seed cultures of *C. glutamicum* were raised at 30 °C and 150 rpm in 2TY medium (containing 16 g/L tryptone, 10 g/L yeast extract, and 5 g/L NaCl). Microscale cultivations were run in a BioLector Pro microbioreactor system (m2p Labs). These experiments used 48-well FlowerPlates covered with gas-permeable sealing membranes (Beckman Coulter), with a working volume of 1 mL per well, operated at 30 °C and 1100 rpm. Shake-flask fermentations were carried out with a working volume of 50 mL at 30 °C and 150 rpm using baffled flasks. The formulation for the minimal medium CGXII followed established protocols [23], with additions of 0.2 mg/L biotin and 1 mL/L trace element solution. Specifically, the base CGXII salt solution consisted of 10 g/L (NH₄)₂SO₄, 1 g/L KH₂PO₄, 1 g/L K₂HPO₄, 5 g/L urea, 42 g/L MOPS, alongside 1 mL/L of a magnesium stock (250 g/L MgSO₄·7H₂O) and 1 mL/L of a calcium stock (13.25 g/L CaCl₂·2H₂O). The accompanying trace metal stock solution contained 16.4 g/L FeSO₄·7H₂O, 10 g/L MnSO₄·H₂O, 1 g/L ZnSO₄·7H₂O, 0.31 g/L CuSO₄·5H₂O, and 0.02 g/L NiCl₂·6H₂O, with the pH brought down to 1 using HCl. Cultures were started at an initial target OD₆₀₀ of 1 approx. Cell growth from flask cultivations was monitored via optical density measurements on a Biochrom Ultrospec 7500 spectrophotometer (Fisher Scientific). To calculate the dry biomass concentration (g/L), the formula 0.343 × (OD₆₀₀ final) was applied, where 0.343 represents the established local correlation factor for *C. glutamicum* [24]. The overall biomass yield was determined as the mass of generated cells per unit mass of depleted carbon substrate. Where necessary, cultivation media were supplemented with selective antibiotics or inducers at final concentrations of 1 mM IPTG, or 5 µg/mL tetracycline. Basal media components were sterilized via autoclaving (121 °C for 20 min). Heat-sensitive additives, including the trace metal mix, biotin, IPTG, and antibiotics, were sterilized independently using 0.22 µm syringe filters.

**Table 1:** strains and plasmids used in this study.

| Stain/Plasmid | Description | Reference |
| --- | --- | --- |
| Strains |  |  |
| *E. coli* DH5α | Cloning host used for plasmid propagation and screening. Genotype: $\Delta lacU169$ ($\phi 80 lacZ$ $\Delta M15$), *supE44*, *hsdR17*, *recA1*, *endA1*, *gyrA96*, *thi-1*, *relA1*. | [25] |
| *C. glutamicum* | Wild-type strain ATCC 13032, auxotrophic for biotin. | [26] |
| DM1729SL | Lysine producer. *C. glutamicum* ATCC13032 with the following modifications: *pyc*<sup>P458S</sup>, *hom*<sup>V59A</sup>, *lysC*<sup>T311I</sup>, $\Delta sugR$, and $\Delta ldhA$. Also known as DM1729$\Delta sugR\Delta ldhA$. | [17] |
| Plasmids |  |  |
| pECTX99a | Tet<sup>R</sup>, *C. glutamicum*/*E. coli* shuttle vector containing *Ptrc*, *lacIq*, and pGA1 oriVCg | [27] |
| pECTX99a-*ectABCD-ask* | pECTX99a derivative carrying *ectABCD* and *ask* from *P. stutzeri* DSM 5190 | This work |
| pECTX99a-hectX | pECTX99a-based combinatorial library containing the *ask-ectABCD* operon under the control of diverse pTIR combinations | This work |
| pECTX99a-hect46 | pECTX99a derivative harboring the *ask-ectABCD* operon with optimized pTIR configuration for hydroxyectoine production | This work |

### 2.2 Molecular genetic techniques, combinatorial assembly, and strains construction

Routine recombinant DNA protocols were carried out following standard methodologies [28]. Cloning-grade PCR were conducted using CloneAmp™ HiFi PCR Premix (Takara Bio Inc.), while colony screening via PCR was executed with GoTaq® DNA Polymerase (Promega). *E. coli* DH5α served as the host for cloning, with transformation performed via heat-shock [28]. For *C. glutamicum* strains, transformations were performed via electroporation using Elepo21 (Nepa gene) with chilled cuvettes and the following settings: 1 poring pulse at 2 kV, 3.5 ms pulse length, 50 ms pulse interval, + polarity, and 3 transfer pulses with 0.2 kV, 50 ms pulse length, 50 ms pulse interval, and +/-polarity. After electroporation, the cells were subjected to a heat shock at 46 °C for 5 min and 45 s, followed by recovery in 2YT medium for 1 h at 30 °C. The cells were then plated on selective agar and incubated overnight at 30 °C.

All hydroxyectoine biosynthesis genes (*ectABCD-ask*) were amplified from the genomic DNA of *P. stutzeri* DSM 5190. The complete *ectABCD-ask* cluster was amplified using the primer pair HectFw and HectRv. To introduce different pTIRs through combinations of ribosome binding sites (RBSs) and translational initiation sites (TISs), the individual genes were amplified using specific forward primers paired with a common reverse primer: *ask* via forward primers K(AA)Fw, K(AG)Fw, or K(TG)Fw combined with K(XX)Rv; *ectAB* via AB(AA)Fw, AB(AG)Fw, or AB(TG)Fw combined with AB(XX)Rv; *ectC* via C(AA)Fw, C(AG)Fw, or C(TG)Fw combined with C(XX)Rv; and *ectD* via D(AA)Fw, D(AG)Fw, or D(TG)Fw combined with D(XX)Rv. pTIR values were predicted utilizing the Salis Lab RBS Calculator online platform (Table 2)[29]. Transformant screening was performed by colony PCR using the primer ECFw/ECRv. Details regarding primer designations, sequences, and applications can be found in Table S1. The base vectors pECXT99a [27] underwent restriction digestion with BamHI, followed by a Gibson Assembly reaction [30] to integrate the corresponding purified PCR amplicons.

**Table 2:**
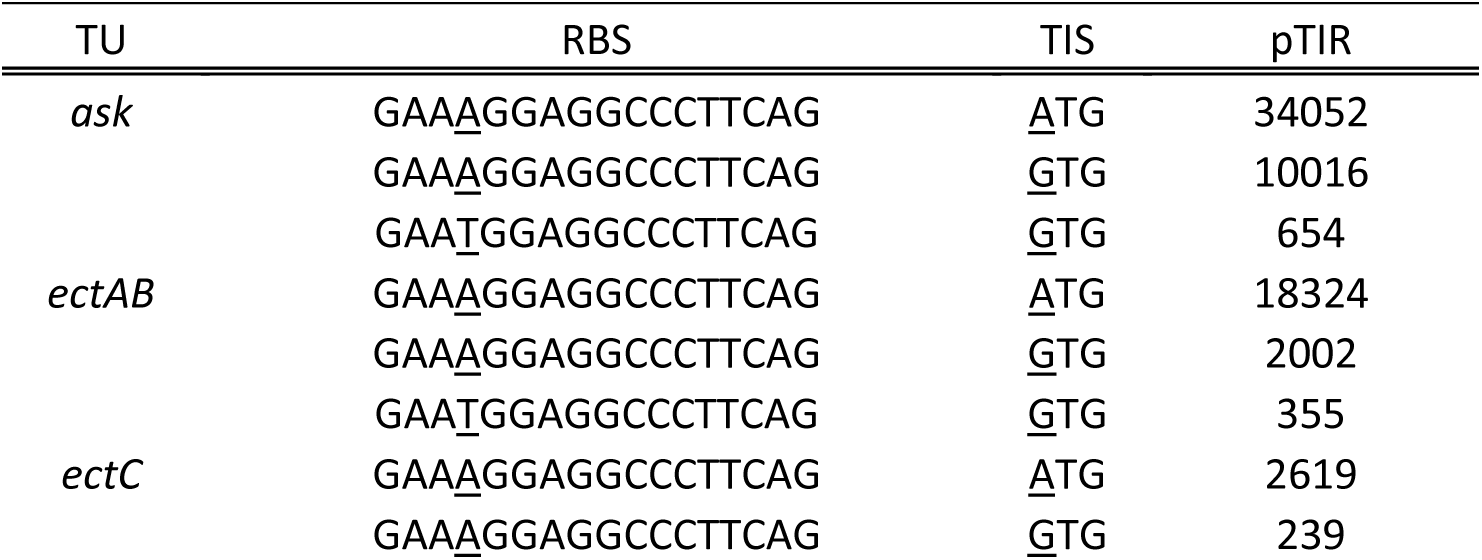
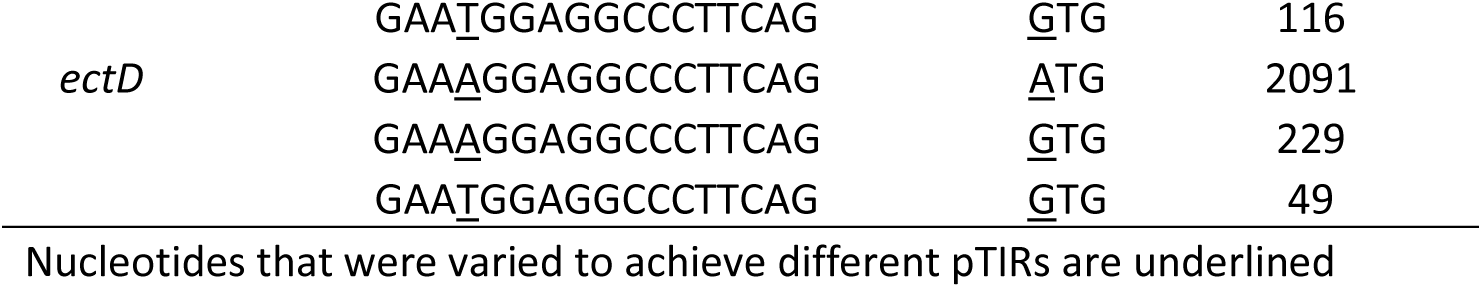
pTIR values per TU by combining different RBS and TIS sequences.

### 2.3 Combinatorial library assembly and screening

To construct the hydroxyectoine expression library, the BamHI-linearized pECXT99a vector was blended with the heterogeneous *ask*, *ectAB*, *ectC*, and *ectD* PCR products featuring randomized RBS and TIS regions. This reaction utilized equimolar mass proportions of 120 ng per DNA component within a standard Gibson Assembly master mix. The assembly reaction was kept at 50 °C for 1 h [30], and a 10 μl aliquot of the mixture was subsequently transformed into competent *E. coli* DH5α cells. 400 positive *E. coli* recombinant colonies were isolated, scraped and pooled into a single batch to isolate the collective plasmid library named pECXT99a-hectX. The purified plasmid library was then introduced into *C. glutamicum* wild type via electroporation. Transformants were recovered on 2YT agar plates supplemented with 5 μg/mL tetracycline. *C. glutamicum* transformants were subsequently growth evaluated in a BioLector Pro micro-fermentation system (m2p-labs) at 40°C using Flower Plates filled with 1% glucose 2TY, alongside the control strain *C. glutamicum* pECXT99a-*ectABCD-ask*.

### 2.4 HPLC analyses

High-performance liquid chromatography (HPLC) (Waters Alliance e2695) was used for the quantification of extracellular ectoine, hydroxyectoine, lysine, alanine, glucose, trehalose, and lactate in cell culture supernatants. To collect supernatants, 1 mL of culture was transferred into a microcentrifuge tube and centrifuged at 17,000xg for 15 min to remove cells. The supernatant was then transferred to a new tube and frozen at-20°C until use.

For the quantification of ectoine and hydroxyectoine, supernatants were diluted 1:10 using 75:25 Acetonitrile:Water. Separation was carried out on a Luna Silica Column (250 x 4.6 mm, 5 µm, Phenomex) at 30 °C. Detection was performed via a 2489 UV detector at 210 nm (Waters). The mobile phases were buffer A 10 mM monopotassium phosphate pH 6.0, and buffer B 100% HPLC-grade acetonitrile, applied at the flow rate of 1 mL/min with the isocratic run blend of 15% buffer A / 85% Buffer B. For the quantification of amino acids lysine and alanine, 1:20 diluted culture supernatants were derivatized with fluorenylmethoxycarbonyl chloride (FMOC) as described previously [31]. Separation was carried out on a Symmetry C18 column (125 × 4.6 mm, 3.5 μm; Waters) at 25 °C. Detection was performed via a 2475 fluorescence detector (Waters). The mobile phases were buffer A (50 mM sodium acetate, pH 4.2) and buffer B (100% acetonitrile), applied at the flow rate of 1.3 mL/min with the following gradient: 0 min, 62% A / 38% B; 5 min, 62% A / 38% B; 12 min, 43% A / 57% B; 14 min, 24% A / 76% B; 15 min, 43% A / 57% B; and 18 min, 62% A / 38% B. For carbohydrates metabolites HPLC analysis, culture supernatants were diluted to 1:10. Glucose, trehalose, and lactate were quantified using an Aminex HPX-87H column (300mm x 7.8mm, Bio-Rad) at 60 °C and detected by a refractive index 2414 RI detector (Waters). Sulfuric acid 5 mM was used as mobile phase at 0.6 mL/min.

### 2.5 Bioprocesses in lab-bench bioreactors

Controlled cultivations were performed in 1.25 L baffled glass vessels (Applikon Biotechnology) fitted with twin 45 mm Rushton impellers fixed at heights of 6 and 12 cm from the bottom of the vessel. Online tracking of pH and relative dissolved oxygen (rDO) was handled via 12 mm AppliSens electrochemical probes. The culture pH was kept constant at 7.0 through automated additions of 10% (w/w) phosphoric acid and 4 M potassium hydroxide. To sustain the target rDOS at either 30% or 50%, an automated feedback loop adjusted the agitation intensity within a safe operational window of 200 to 800 rpm. Foaming was managed by manual additions of Antifoam 204 as needed. A constant airflow of 0.75 vvm was supplied through an L-shaped sparger, and an integrated heating jacket regulated the process temperature at 30 °C or 40 °C. The starting volume for the batch stage was set to 0.8 L. Process vessels were inoculated using overnight seed cultures raised in 2TY medium containing 1% glucose. The baseline batch cultivation medium was formulated to contain the following components per liter: 20 g (NH_4_)_2_SO_4_, 10 g urea, 0.8 g yeast extract, 0.5 g KH_2_PO_4_, 0.5 g K_2_HPO_4_, 0.01325 g CaCl_2_·2H_2_O, 0.25 g MgSO_4_·7H_2_O, 0.2 mg biotin, 0.005 g tetracycline, 0.083 g IPTG, and 1 mL trace element solution consisting of FeSO_4_·7H_2_O (0.025 g/L), MnSO_4_·H_2_O (0.015 g/L), ZnSO_4_·7H_2_O (0.025 g/L), CuSO_4_ (0.05 mg/L), and NiCl_2_·6H_2_O (0.005 mg/L). 5% glucose was added as carbon source during the batch phase. Additionally, 20mL of 2YT were added to the batch medium. During carbon-limited fed-batch cultivations, the concentrated feed medium consisted of 200 g/L glucose, 0.2 mg/L biotin, 0.005 g/L tetracycline, 0.083 g/L IPTG, 1 g/L yeast extract, and 1 mL/L of the trace element formulation. To avoid the accumulation of residual glucose in the vessel, the volumetric feed rate was modulated manually within a range of 0.137 to 0.096 mL/min.

## 3. Results

### 3.1. Hydroxyectoine enhances the growth of *C. glutamicum* under heat stress

Compatible solutes such as ectoine and hydroxyectoine contribute to cellular protection under environmental stress by stabilizing proteins, membranes, and other macromolecular structures. Hydroxyectoine has been particularly associated with enhanced protection against elevated temperatures [21]. Therefore, the effect of growth temperature on *C. glutamicum* was first assessed, followed by an evaluation of whether supplementation with ectoine and hydroxyectoine could alleviate heat-induced growth impairment.

*C. glutamicum* (pECXT99a) was grown in 2YT medium containing 1% glucose at temperatures ranging from 30 to 42.5 °C. Under these conditions, no significant differences in biomass formation or growth rate were observed at 30, 35, or 37.5 °C. However, growth impairment became evident from 40 °C (Fig. 2A). To evaluate the potential roles of ectoine and hydroxyectoine as thermoprotectants, *C. glutamicum* (pECXT99a) was subsequently grown in 2YT medium containing 1% glucose and supplemented with either 5 mM ectoine, 5 mM hydroxyectoine, a combination of both compounds at 5 mM each, or no supplementation. The experiments were performed at 30 °C (Fig. 2B) and 40 °C (Fig. 2C). At 30 °C, no significant differences in biomass formation or growth rate were observed among the tested conditions. However, when *C. glutamicum* (pECXT99a) was grown at 40 °C, supplementation with 5 mM hydroxyectoine increased both biomass formation and growth rate by approximately 20%. Interestingly, simultaneous supplementation with ectoine and hydroxyectoine did not produce the same positive effect (Fig. 2C). This finding was further explored in the subsequent section of this study.

**Fig. 2:**
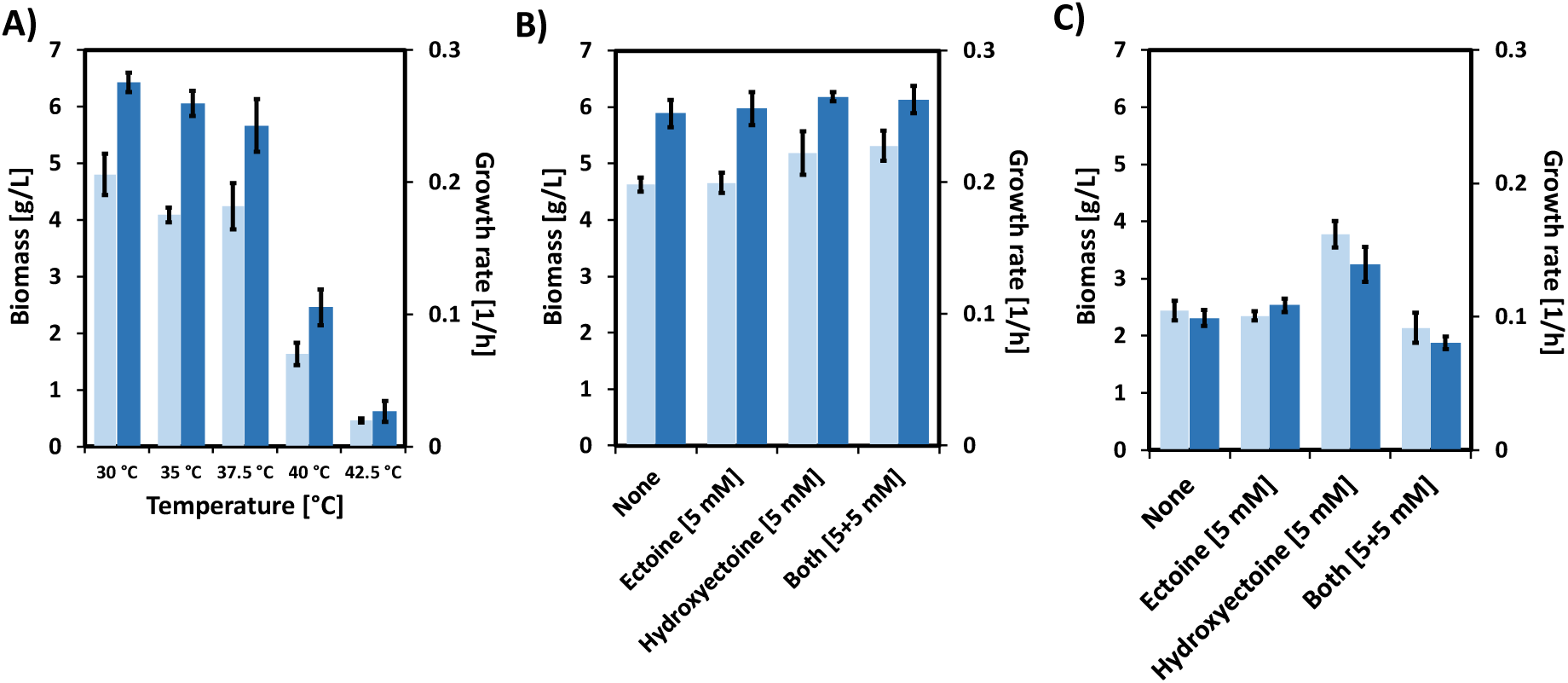
Effect of temperature and compatible-solute supplementation on the growth of *C. glutamicum*. **(A)** Biomass formation and specific growth rate during cultivation at temperatures ranging from 30 to 42.5 °C. **(B-C)** Effect of supplementation with 5 mM ectoine, 5 mM hydroxyectoine, or both compounds at 5 mM each on biomass formation and growth rate at 30 °C **(B)** and 40 °C **(C)**. Cultures without compatible-solute supplementation were used as controls. Light-blue bars represent final biomass concentrations, whereas dark-blue bars represent specific growth rates. Error bars indicate the variability among replicates.

### 3.2 Thermal stress-based screening of a synthetic hydroxyectoine operon

The ability of hydroxyectoine to partially alleviate growth impairment in *C. glutamicum* under non-optimal temperature conditions was exploited in this study as a selective pressure for the construction of a randomly assembled synthetic hydroxyectoine operon aimed at achieving efficient hydroxyectoine production. As detailed in the Materials and Methods, the synthetic operon was designed to contain four translational units (*ask*, *ectAB*, *ectC*, and *ectD*) with three possible pTIRs for each unit (Fig. 3A). Consequently, random assembly of the four units in the fixed order *ask-ectAB-ectC-ectD* could generate 81 possible operon variants. The combinatorial plasmid library was transformed into *C. glutamicum*. Subsequently, 174 *C. glutamicum* transformants, designated *C. glutamicum* (pECXT99a-*hectX*), were screened at 40 °C in BioLector FlowerPlates containing 1 mL of 2YT medium supplemented with 1% glucose. The strain *C. glutamicum* (pECXT99a-*ectABCD-ask*), which overexpressed the pathway genes in their native operon configuration, was used as the control. Growth rates and biomass values were calculated for the 174 *C. glutamicum* transformants and the control strain (Fig. 3B). Under these conditions, the control strain exhibited a growth rate of 0.04 ± 0.00 1/h and reached a final biomass concentration of 1.3 ± 0.0 g/L (Fig. 3B). In contrast, the *C. glutamicum* (pECXT99a-*hectX*) transformants displayed diverse growth profiles. Seven transformants, 46, 82, 86, 112, 128, 131, and 138, showed growth rates above 0.1 1/h and final biomass concentrations above 2 g/L (Fig. 3B). Ectoine and hydroxyectoine production by these seven transformants and the control strain was subsequently quantified by HPLC (Fig. 3C). The control strain produced 3.1 mM ectoine and 1.8 mM hydroxyectoine, corresponding to a hydroxyectoine-to-ectoine ratio of 0.6. By comparison, the seven selected transformants produced 0.8-1.1 mM ectoine and 3.6-4.7 mM hydroxyectoine. Transformant 46 exhibited the highest hydroxyectoine-to-ectoine ratio, reaching a value of 4.7 (Fig. 3C). Additionally, four transformants, 7, 61, 101, and 149, displaying low growth rates were randomly selected (Fig. 3C). Their ectoine and hydroxyectoine titers were also quantified, revealing production values of 1.8-2.0 mM hydroxyectoine and 2.5-2.8 mM ectoine, yielding hydroxyectoine-to-ectoine ratios of 0.7. Sequencing of the plasmids carried by the fast-and slow-growing transformants revealed the assembled TUs and, consequently, their corresponding pTIR values (Table S2). The pTIR assigned to *ask* was consistently the highest available value in the selected transformants 46, 82, 86, 112, 128, 131, and 138, whereas the control strain carried a lower *ask* pTIR. The opposite pattern was observed for *ectAB*, for which the control strain displayed a higher pTIR than the selected transformants. For *ectC*, the pTIRs of the selected transformants ranged from 116 to 239 a.u., compared with 961 a.u. in the control strain. Finally, *ectD* exhibited a pTIR of 2091 a.u. in all selected transformants, whereas the corresponding value in the control strain was 1237 a.u. (Fig. 3D, Table S2). On the other hand, transformants 7, 61, 101, and 149 showed pTIR values of 10016, 18324, 2619, and 229 a.u. for *ask*, *ectAB*, *ectC*, and *ectD*, respectively (Fig. 3D, Table S2). Selected plasmid from transformant 46, designated pHect46, was chosen for further characterization in this study.

**Fig. 3:**
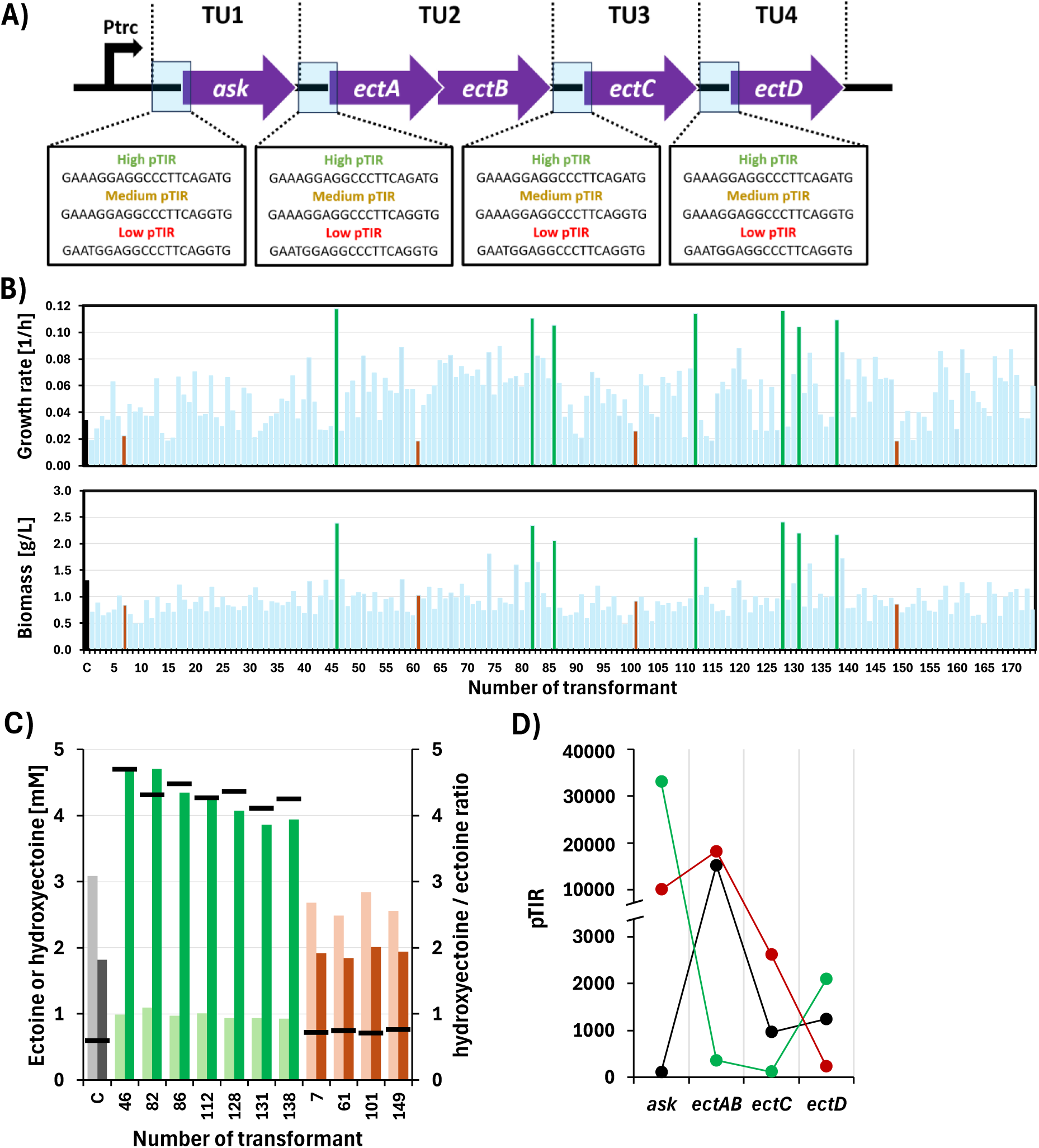
Thermal stress-based screening of a combinatorial synthetic hydroxyectoine operon library in *C. glutamicum*. **(A)** Schematic representation of the synthetic operon containing four translational units, TU1 (*ask*), TU2 (*ectAB*), TU3 (*ectC*), and TU4 (*ectD*), each assembled with one of three possible predicted translation initiation rates (pTIRs). The resulting combinatorial design generated 81 possible operon configurations. **(B)** Growth rates and final biomass concentrations of 174 *C. glutamicum* transformants (blue bars) screened at 40 °C. The strain carrying the pathway in its native operon configuration was used as the control (C; black bar). Fast-growing transformants selected for further characterization are shown in green, whereas randomly selected slow-growing transformants are shown in red. **(C)** Extracellular ectoine and hydroxyectoine concentrations produced by the control, seven fast-growing transformants, and four slow-growing transformants. Light bars indicate ectoine concentrations, dark bars indicate hydroxyectoine concentrations, and horizontal black lines represent the corresponding hydroxyectoine-to-ectoine ratios. **(D)** pTIR profiles of the four translational units in the control strain (black), representative fast-growing transformant 46, designated CgHect46 (green), and the slow-growing transformants (red).

### 3.3 Improved hydroxyectoine production in a lysine-producing *C. glutamicum* host

In this block of the study, the newly developed plasmid pHect46 was further explored for the production of hydroxyectoine at 30 and 40 °C in flask cultivations. Additionally, hydroxyectoine precursor supply was enhanced by transferring pHect46 into the *C. glutamicum* lysine producer DM1729SL. Hence, minimal medium supplemented with 1% glucose was used to test the strains *C. glutamicum*(pECXT99a) (Fig. 4A), DM1729SL(pECXT99a) (Fig. 4C), *C. glutamicum*(pHect46) (Fig. 4B), and DM1729SL(pHect46) (Fig. 4D) at 30 and 40 °C. Under these conditions, all strains grew well at 30 °C; however, growth was severely impaired at 40 °C, even for the strains harboring plasmid pHect46, contradicting our earlier findings. Two major variables differed from the initial BioLector experiments: the cultivation scale and the medium composition. Consequently, a subsequent round of fermentations was conducted under identical conditions but supplemented with 1 g/L yeast extract. Interestingly, this supplementation improved growth at 40 °C across all strains. Notably, *C. glutamicum*(pHect46) and DM1729SL(pHect46) exhibited growth profiles comparable to those observed at 30 °C (Fig. 4). These results suggest that yeast extract contains specific components that alleviate thermal stress at non-optimal temperatures and/or enhance hydroxyectoine biosynthesis.

**Fig. 4:**
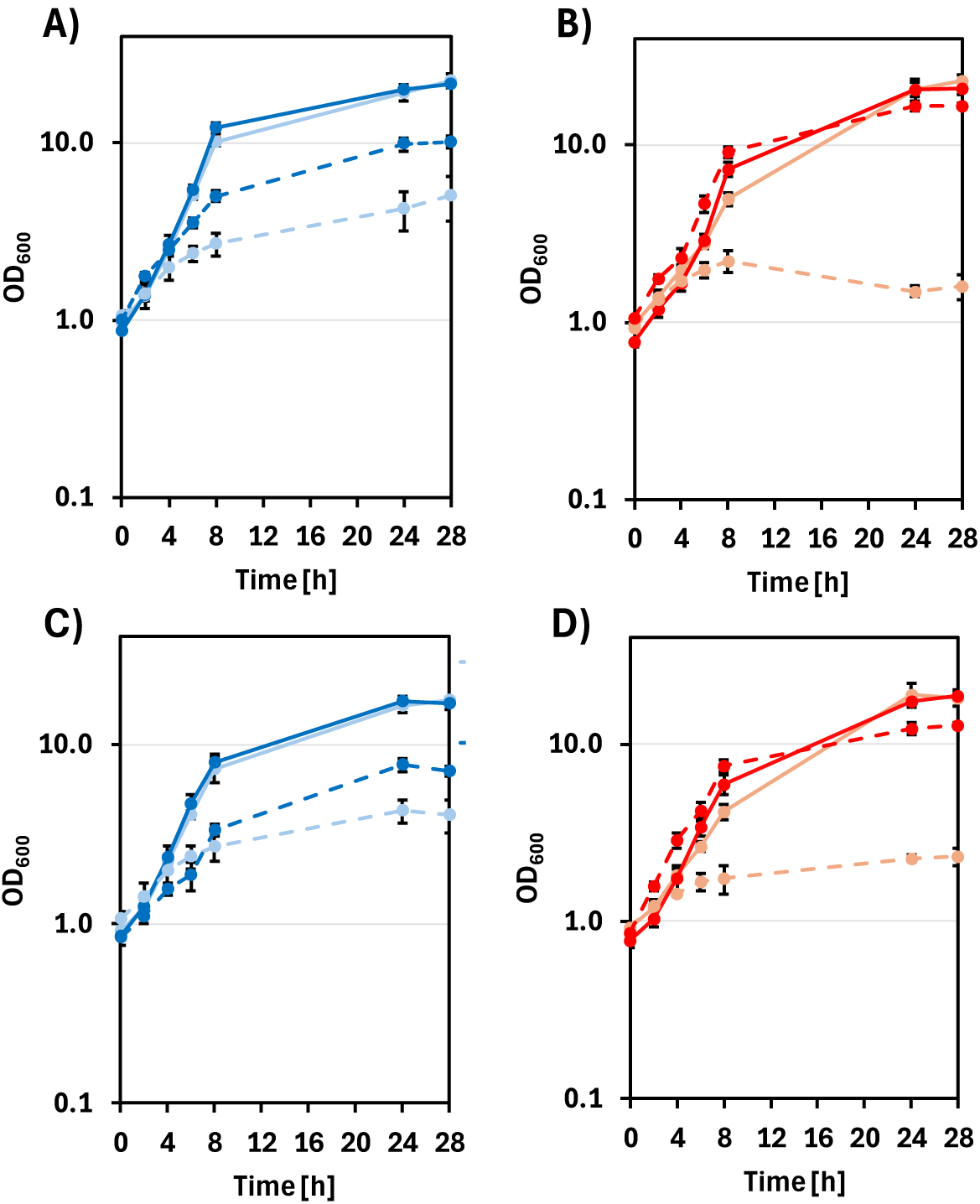
Effects of temperature and yeast extract supplementation on the growth of hydroxyectoine-producing *C. glutamicum* strains. Growth profiles of wild-type *C. glutamicum* carrying (A) pECXT99a or (B) pHect46, and of the lysine-producing strain DM1729SL carrying **(C)** pECXT99a or **(D)** pHect46. Cultivations were performed at 30 °C (solid lines) or 40 °C (dashed lines) in medium without yeast extract supplementation (light colors) or supplemented with yeast extract (dark colors). Data is presented as OD600 over time, and error bars indicate the variability among replicates.

Extracellular titers of lysine, ectoine, and hydroxyectoine were quantified from culture supernatants (Table 3). As expected, the negative control strains *C. glutamicum*(pECXT99a) and DM1729SL(pECXT99a) did not produce ectoine or hydroxyectoine. At 30 °C, strain DM1729SL(pECXT99a) accumulated 7.5 ± 0.4 mM lysine in standard CGXII medium and 7.3 ± 0.5 mM upon yeast extract supplementation. Under identical thermal conditions, *C. glutamicum*(pHect46) yielded 2.0 ± 0.3 mM lysine, 2.0 ± 0.1 mM ectoine, and 4.0 ± 0.3 mM hydroxyectoine. Yeast extract addition slightly shifted these titers to 1.5 ± 0.2, 1.1 ± 0.2, and 4.6 ± 0.4 mM, respectively, though these changes were not statistically significant. In general, higher titers were reached by the strain DM1729SL(pHect46) at 30 °C, with no significant differences regardless of yeast extract supplementation. At 40 °C under yeast extract supplementation, *C. glutamicum*(pHect46) and DM1729SL(pHect46) accumulated 4.7 ± 0.2 and 9.8 ± 0.4 mM hydroxyectoine, respectively. Nevertheless, the maximum hydroxyectoine titer obtained was achieved by strain DM1729SL(pHect46) at 30 °C when supplemented with yeast extract, yielding 11.2 ± 0.6 mM.

**Table 3:** Extracellular lysine, ectoine, and hydroxyectoine concentrations obtained under different flask-cultivation conditions.

| Strain | Temperature | YE | Lysine [mM] | Ectoine [mM] | Hydroxyectoine [mM] |
| --- | --- | --- | --- | --- | --- |
| *C. glutamicum*(pECTX99a) | 30°C | - | 0.0 ± 0.0 | n.d. | n.d. |
|  |  | + | 0.2 ± 0.0 | n.d. | n.d. |
|  | 40°C | - | 0.0 ± 0.0 | n.d. | n.d. |
|  |  | + | 0.0 ± 0.0 | n.d. | n.d. |
| *C. glutamicum*(pHect46) | 30°C | - | 2.0 ± 0.3 | 2.0 ± 0.1 | 4.0 ± 0.3 |
|  |  | + | 1.5 ± 0.2 | 1.1 ± 0.2 | 4.6 ± 0.4 |
|  | 40°C | - | 0.5 ± 0.3 | 0.3 ± 0.1 | 0.4 ± 0.2 |
|  |  | + | 1.4 ± 0.3 | 1.3 ± 0.2 | 4.7 ± 0.2 |
| DM1729SL(pECTX99a) | 30°C | - | 7.5 ± 0.4 | n.d. | n.d. |
|  |  | + | 7.3 ± 0.5 | n.d. | n.d. |
|  | 40°C | - | 0.8 ± 0.2 | n.d. | n.d. |
|  |  | + | 1.1 ± 0.4 | n.d. | n.d. |
| DM1729SL(pHect46) | 30°C | - | 3.4 ± 0.6 | 3.2 ± 0.4 | 10.9 ± 0.9 |
|  |  | + | 3.1 ± 0.4 | 3.1 ± 0.5 | 11.2 ± 0.6 |
|  | 40°C | - | 1.5 ± 0.2 | 2.3 ± 0.0 | 2.3 ± 0.1 |
|  |  | + | 3.1 ± 0.3 | 2.8 ± 0.2 | 9.8 ± 0.4 |

Values are presented as mean ± standard deviation. YE, yeast extract; n.d., not detected.

### 3.4 Effect of temperature and dissolved oxygen on hydroxyectoine production in fed-batch bioreactors

Four carbon-limited fed-batch cultivations were performed in bioreactors to evaluate hydroxyectoine production by strain DM1729SL(pHect46) under controlled conditions (Fig. 5). The cultivations combined two temperatures, 30 and 40 °C, with two rDO setpoints, 30% and 50%. The two rDO setpoints were included because EctD is an ectoine dioxygenase that requires O₂ as a cofactor. Biomass formation, glucose consumption, hydroxyectoine, and by-products formation were monitored over time. Cultivations were terminated when the rDO showed saturation after the feeding indicating carbon source depletion (data not shown).

**Fig. 5:**
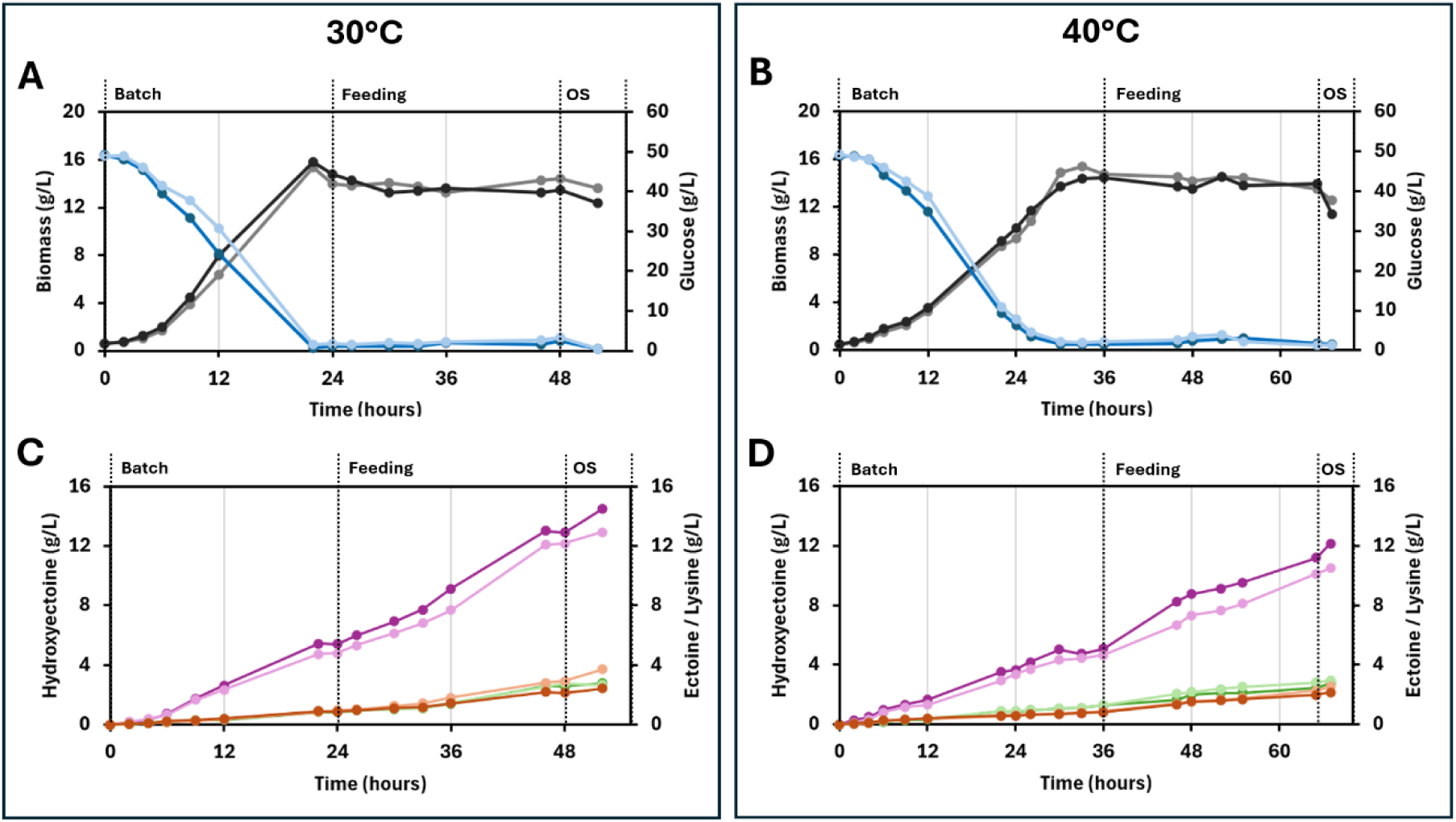
Fed-batch production of hydroxyectoine by DM1729SL(pHect46) under different temperature and dissolved-oxygen conditions. (A, **C)** Cultivations performed at 30 °C and (B, D) cultivations performed at 40 °C. Biomass and glucose concentrations are shown in panels A and B, whereas hydroxyectoine, ectoine, and lysine concentrations are shown in panels C and D. Black lines represent biomass, blue lines glucose, purple lines hydroxyectoine, red lines ectoine, and green lines lysine. Darker shades indicate processes operated at 50% rDO, whereas lighter shades indicate 30% rDO. Vertical dotted lines indicate the transition from batch to feeding phase and the time point at which the osmotic downshock (OS) was applied.

Under these conditions, strain DM1729SL(pHect46) grew faster at 30 °C than at 40 °C, regardless of the rDO setpoint (Fig. 5). In the cultivations performed at 30 °C, the batch phase, initiated with 5% glucose, lasted for 22 h. A linear feed was then started at a rate of 8 mL/h, maintaining the glucose concentration below 0.5 g/L throughout the feeding phases (Fig. 5A). The cultivations were terminated at 48 hours when 200 mL of feeding medium had been pumped into the vessel and the rDO signal showed saturation. Final hydroxyectoine titers of 12.2 and 12.9 g/L were achieved at rDO setpoints of 30% and 50%, respectively. These values corresponded to yield and volumetric productivity values of 0.153 g/g and 0.25 g/L/h at 30% rDO, and 0.161 g/g and 0.26 g/L/h at 50% rDO (Fig. 5C). In contrast, the cultivations performed at 40 °C lasted longer and they were, in general, less productive with regard Hydroxyectoine. Batch phases lasted 33 hours, while feeding phases lasted 32 hours for a total process of 65 hours (Fig. 5B). Hydroxyectoine titers reached 10.1 and 11.2 g/L at rDO setpoints of 30% and 50%, respectively, which corresponded to yields and volumetric productivities of 0.126 g/g and 0.16 g/L/h at 30% rDO, and 0.140 g/g and 0.17 g/L/h at 50% rDO (Fig. 5D). Within the fed-batch cultivations, lysine and ectoine accumulated as by-products under all tested conditions, although their concentrations varied. At 30 °C and 30% rDO, lysine and ectoine reached concentrations of 2.9 and 2.8 g/L, respectively. At 30 °C and 50% rDO, ectoine accumulation was slightly lower (2.1 g/L), possibly reflecting its more efficient conversion to hydroxyectoine (Fig. 5C). Cultivation at 40 °C resulted in ectoine and lysine accumulation of 2.3 and 2.8 g/L at 30% rDO as well as 2.0 and 2.4 g/L at 50% rDO (Fig. 5D).

Following glucose depletion, an osmotic downshock was applied to promote the release of the remaining intracellular hydroxyectoine, following the concept of bacterial milking established by Sauer and Galinski [32]. The reactors were bled to reduce the working volume to 0.5 L. Subsequently, 0.5 L of 10 mM potassium phosphate buffer pH 7.0 was pumped into the reactors, restoring the final volume to 1 L. Two hours later, samples were collected for biomass determination and HPLC analysis. After correction for the twofold dilution introduced by the downshock procedure, extracellular hydroxyectoine titers increased by 4-6% in cultivations operated at 30% rDO and by 8-12% in cultivations operated at 50% rDO. Overall, the best production performance was obtained at 30 °C and 50% rDO, reaching 12.9 g/L hydroxyectoine after glucose depletion and 14.5 g/L after osmotic downshock (Fig. 5C and D).

## 4. Discussion

Microorganisms exposed to osmotic, thermal, desiccation, and other environmental stresses frequently accumulate compatible solutes to preserve cellular function without interfering with central metabolism [1]. Among these compounds, ectoine and hydroxyectoine are cyclic amino acid derivatives produced by diverse bacteria and some archaea [1]. Hydroxyectoine is a high-value extremolyte with potential applications in cosmetics, pharmaceuticals, biotechnology, diagnostics, and biological-material preservation [1,4,33]. Its ability to stabilize proteins, membranes, nucleic acids, and whole cells under thermal, osmotic, freezing, and desiccation stress makes it particularly attractive as a protective ingredient and formulation excipient [3]. Hydroxyectoine has been shown to reduce protein unfolding and aggregation, preserve enzyme activity under adverse conditions, stabilize therapeutic antibodies during spray drying and freeze [34]. These properties support its potential use in biopharmaceutical formulations, industrial enzyme preparations, diagnostic devices, and cryopreservation systems. Hydroxyectoine has also been investigated for cytoprotective and anti-aggregation effects relevant to hypoxic injury and neurodegenerative disorders [35]. In cosmetics, ectoine-family compounds are used for hydration, barrier support, and protection against environmental stress, although most clinical and commercial evidence currently relates to ectoine rather than hydroxyectoine specifically [36]. In this study, externally supplemented hydroxyectoine was shown to partially alleviate thermal stress in *C. glutamicum* (*Fig. 2*). Interestingly, supplementation with ectoine and hydroxyectoine together did not produce the same effect. Although *C. glutamicum* possesses well-characterized secondary transporters for the compatible solutes ectoine, proline, and betaine, the specific uptake and physiological handling of hydroxyectoine may rely on overlapping substrate specificities within these compatible-solute transport systems or on transport activities identified in engineered whole-cell biotransformation systems [19,37]. It is therefore possible that ectoine and hydroxyectoine share and compete for the same uptake systems, although, to our knowledge, this has not yet been reported.

Hydroxyectoine is currently obtained predominantly through microbial processes. Traditional production relies on natural halophilic bacteria cultivated under high-salinity conditions that stimulate compatible-solute biosynthesis, followed by product recovery from the cells or fermentation broth [12]. However, the use of highly saline media causes equipment corrosion and complicates downstream processing and wastewater treatment [12]. Consequently, engineered non-halophilic microorganisms have emerged as promising alternatives for hydroxyectoine production because they can operate without the high salt concentrations required by many natural producers. Among these hosts, *E. coli* has been extensively engineered for *de novo* synthesis from glucose. In particular, optimization of the expression levels of the ectoine and hydroxyectoine biosynthetic genes from *Halomonas elongata*, combined with dynamic regulation of the intracellular 2-oxoglutarate pool, enabled strain HECT31 to produce 14.9 g/L hydroxyectoine without detectable ectoine accumulation or osmotic-stress induction [38]. More recently, further metabolic engineering of *E. coli* included screening of ectoine hydroxylases, improvement of L-2,4-diaminobutyrate formation, activation of the glyoxylate cycle, and balancing of 2-oxoglutarate distribution. The optimized strain produced 3.4 g/L hydroxyectoine in shake flasks and reached 58 g/L through a semi-continuous feeding process in NaCl-free medium [39]. *C. glutamicum* is another attractive host because of its established industrial use, robustness, and naturally high metabolic flux through the aspartate-family amino acid pathway [40]. This pathway supplies L-aspartate-β-semialdehyde, the direct precursor used by EctB for ectoine biosynthesis. Initial *de novo* production studies with recombinant *C. glutamicum* expressing *ectABCD* from *P. stutzeri* demonstrated simultaneous ectoine and hydroxyectoine formation, although hydroxyectoine titers remained low, at approximately 0.4 g/L [13]. A different strategy subsequently exploited *C. glutamicum* as a whole-cell catalyst for the hydroxylation of externally supplied ectoine [19]. By screening heterologous EctD enzymes and optimizing cultivation conditions, a strain expressing the *ectD* gene from *P. stutzeri* produced 74 g/L hydroxyectoine with approximately 70% selectivity [19]. The same study also demonstrated a two-stage process in which ectoine was first produced fermentatively by one engineered *C. glutamicum* strain and subsequently converted into hydroxyectoine by an EctD-expressing strain without intermediate ectoine purification [19].

Establishing efficient single-strain *de novo* production in *C. glutamicum* therefore remains particularly attractive, as it could combine the host’s industrial robustness and strong aspartate-family metabolism with a simpler process based directly on conventional carbon substrates. In the present study, this challenge was addressed by combinatorially balancing precursor supply, ectoine biosynthesis, and ectoine hydroxylation in a single engineered *C. glutamicum* strain. In a previous riboflavin research, the pTIRs of the *ribGCAH* genes were varied to generate synthetic operons with different relative expression profiles. Screening of these constructs identified an operon architecture that increased riboflavin production without impairing growth, demonstrating that pathway performance depended on balanced rather than uniformly strong expression of all biosynthetic genes [41]. Combinatorial adjustment of translation initiation has previously been used to explore multidimensional enzyme-expression spaces. Zelcbuch et al. combined a compact set of characterized RBSs with multiple genes to span a high-dimensional expression space and identify combinations that balanced protein levels. Their method similarly recognized that optimal pathway performance cannot generally be achieved by maximizing the expression of every component [42]. Farasat et al. extended this concept through computationally designed degenerate RBS libraries that sampled broad ranges of predicted translation rates in several bacterial hosts [43]. Jeschek et al. subsequently developed the RedLibs strategy to design rationally reduced RBS libraries that uniformly cover a desired translation-initiation space while limiting library size. They demonstrated the approach by balancing the violacein pathway around a metabolic branch point [20]. Other studies have explored broader forms of combinatorial pathway refactoring.

Nowroozi et al. combined RBS variants with modular gene assembly to optimize multienzyme pathways, while Smanski et al. varied gene order, orientation, regulatory elements, and operon organization in refactored nitrogen-fixation gene clusters [44,45]. These strategies investigate a larger architectural space than the present study, in which gene order was fixed and only translational control was varied. Fixing the order of *ask-ectAB-ectC-ectD* in this study substantially reduced library complexity and facilitated interpretation of the selected expression patterns, particularly the recurring combination of strong predicted translation of *ask* and *ectD* with lower predicted translation of *ectAB* and *ectC* (Fig. 3). Three pTIR levels were assigned to *ask*, *ectAB, ectC*, and *ectD*. Rather than screening directly by product quantification, hydroxyectoine-associated thermoprotection was used to identify promising variants. HPLC analysis showed that fast-growing transformants generally produced more hydroxyectoine, accumulated less ectoine, and displayed higher hydroxyectoine-to-ectoine ratios than the native-operon control. Thus, thermal stress provided a useful, though indirect, proxy for pathway performance (Fig. 3). Additionally, it was observed that yeast extract supplementation had a pronounced effect on growth at 40 °C (Fig. 4). Yeast extracts may instead supply amino acids, vitamins, trace nutrients, or other growth factors that support cellular maintenance and stress responses at elevated temperature [46]. At 30 °C, however, yeast extract did not significantly increase hydroxyectoine titers (Table 3), indicating that its main contribution was improved physiological robustness rather than a direct stimulation of product formation, although the component responsible for this effect remains unknown.

Carbon-limited fed-batch cultivation substantially increased hydroxyectoine production by DM1729SL(pHect46), with the best performance obtained at 30 °C and 50% rDO, reaching 14.5 g/L hydroxyectoine after complete glucose depletion and subsequent application of an osmotic downshock. This titer clearly exceeds those reported for several natural or moderately engineered hydroxyectoine producers. For example, fermentation optimization of the natural producer *H. salina* BCRC17875 yielded 2.9 g/L hydroxyectoine, whereas an engineered *H. salifodinae* strain reached 4.9 g/L in fed-batch cultivation after improving *ectD* expression, preventing ectoine degradation, and increasing 2-oxoglutarate supply [10,11]. The osmotic downshock applied here follows the principle of “bacterial milking,” originally developed for *H. elongata*, in which cells grown at high salinity were subjected to repeated hypoosmotic shocks to release intracellular ectoine. Sauer and Galinski reported that this strategy could be repeated for at least nine cycles and achieved an ectoine biomass productivity of about 0.155 mg/g cell dry weight per cycle [32].

Engineered *E. coli* currently provides the strongest comparison for *de novo* hydroxyectoine synthesis. Strain HECT31 produced 14.9 g/L hydroxyectoine without detectable ectoine accumulation by dynamically controlling the intracellular 2-oxoglutarate pool [38]. A more recent *E. coli* fed-batch process reached 8.6 g/L at a productivity of 0.24 g/L/h in a 7-L bioreactor [47]. Another extensively engineered *E. coli* strain recently reached 58 g/L through a semi-continuous feeding process in NaCl-free medium, demonstrating the remaining potential of intensive strain and process optimization [39]. The improved performance at 50% compared with 30% rDO is consistent with the oxygen requirement of EctD [48]. At 30 °C, the higher rDO setpoint increased the hydroxyectoine titer and productivity while slightly reducing ectoine accumulation, suggesting more efficient conversion of ectoine into hydroxyectoine (Fig. 5). Nevertheless, the remaining ectoine indicates that EctD activity continued to limit pathway completion. Because the reaction also requires Fe²⁺ and 2-oxoglutarate, increasing oxygen supply alone may not be sufficient. The high-performing *E. coli* and *H. salifodinae* strains were improved partly by manipulating 2-oxoglutarate availability [10,38], highlighting this metabolite as a logical target for further engineering of DM1729SL(pHect46).

Hydroxyectoine production was also demonstrated in bioreactors operated at 40 °C. Although titers and productivities were lower than those obtained at 30 °C, this result demonstrates the feasibility of production at elevated temperature. Such processes could potentially reduce reactor-cooling requirements, an advantage also discussed for thermophilic production hosts such as *Bacillus methanolicus*, which grows optimally at approximately 50 °C [49].

## 5. Conclusion

Overall, this study establishes *C. glutamicum* as a promising platform for *de novo* hydroxyectoine production. Thermal stress-based combinatorial pathway balancing enabled the identification of pHect46, while enhanced precursor supply and controlled fed-batch cultivation in combination with osmotic downshock substantially improved hydroxyectoine production. The best performance was obtained at 30 °C and 50% rDO, approaching reported *E. coli* benchmarks and surpassing previous *de novo* production in *C. glutamicum*. Remaining ectoine and lysine accumulation indicate clear opportunities to further improve pathway selectivity, EctD activity, and carbon allocation.

## Author Contributions

**Luciana Fernandes Brito**: investigation, writing-original draft, writing-review and editing, methodology, conceptualization. **Nathalie van Assel**: investigation, writing-review and editing. **Fernando Pérez-García**: investigation, writing-original draft, writing-review and editing, supervision, project administration, methodology, validation, conceptualization, funding acquisition.

## Supporting information

Supplementary material

## Acknowledgements

No applicable.

