## Supplementary material for "Thermal stress-based pathway engineering and bioprocess optimization of *Corynebacterium glutamicum* for *de novo* hydroxyectoine production"

**Table S1**: list of primers used in this study

| Primer name | Sequence (5->3) | Description |
|---|---|---|
| ECFw | **TTTGCGCCGACATCATAACG** | Forward primer for colony PCR with the plasmid pECXT99a |
| ECRv | **TACTGCCGCCAGGCAAATTC** | Reverse primer for colony PCR with the plasmid pECXT99a |
| HectFw | **ATGGAATTCGAGCTCGGTACCCGGGGAAAGGAGGCCCTTCAGATGCCTACCCTAAAAAGGAATTC** | Forward primer for the amplification of *ectABCD-ask* from *P. stutzeri* |
| HectRv | **GCCTGCAGGTCGACTCTAGAGGATCTCAGGCCGCGGCAATCACGTCG** | Reverse primer for the amplification of *ectABCD-ask* from *P. stutzeri* |
| K(AA)Fw | **AGACCATGGAATTCGAGCTCGGTACCCGGGGAAAGGAGGCCCTTCAGATGCATACCGTGGAAAAGATCG** | Forward primer 1 for the amplification of *ask* from *P. stutzeri*. High pTIR for *ask* |
| K(AG)Fw | **AGACCATGGAATTCGAGCTCGGTACCCGGGGAAAGGAGGCCCTTCAGGTGCATACCGTGGAAAAGATCG** | Forward primer 2 for the amplification of *ask* from *P. stutzeri*. Medium pTIR for *ask* |
| K(TG)Fw | **AGACCATGGAATTCGAGCTCGGTACCCGGGGAATGGAGGCCCTTCAGGTGCATACCGTGGAAAAGATCG** | Forward primer 3 for the amplification of *ask* from *P. stutzeri*. Low pTIR for ask |
| K(XX)Rv | **TCAGGCCGCGGCAATCACGTCG** | Reverse primer for the amplification of *ask* from *P. stutzeri* |

| | | |
|---|---|---|
| AB(AA)Fw | AACCATGGCGACGTGATTGCCGCGGCCTGAGAAAGGAGGCCCTTCAGATGCCTACCCTAAAAAGGAATTC | Forward primer 1 for the amplification of *ectAB* from *P. stutzeri*. High pTIR for *ectAB* |
| AB(AG)Fw | AACCATGGCGACGTGATTGCCGCGGCCTGAGAAAGGAGGCCCTTCAGGTGCCTACCCTAAAAAGGAATTC | Forward primer 2 for the amplification of *ectAB* from *P. stutzeri*. Medium pTIR for *ectAB* |
| AB(TG)Fw | AACCATGGCGACGTGATTGCCGCGGCCTGAGAATGGAGGCCCTTCAGGTGCCTACCCTAAAAAGGAATTC | Forward primer 3 for the amplification of *ectAB* from *P. stutzeri*. Low pTIR for *ectAB* |
| AB(XX)Rv | TCAGGAAGCTTGGTTCTCGGTC | Reverse primer for the amplification of *ectAB* from *P. stutzeri* |
| C(AA)Fw | AGCGAGCAGACCGAGAACCAAGCTTCCTGAGAAAGGAGGCCCTTCAGATGATCGTCAGAACCCTCGCCG | Forward primer 1 for the amplification of *ectC* from *P. stutzeri*. High pTIR for *ectC* |
| C(AG)Fw | AGCGAGCAGACCGAGAACCAAGCTTCCTGAGAAAGGAGGCCCTTCAGGTGATCGTCAGAACCCTCGCCG | Forward primer 2 for the amplification of *ectC* from *P. stutzeri*. Medium pTIR for *ectC* |
| C(TG)Fw | AGCGAGCAGACCGAGAACCAAGCTTCCTGAGAATGGAGGCCCTTCAGGTGATCGTCAGAACCCTCGCCG | Forward primer 3 for the amplification of *ectC* from *P. stutzeri*. Low pTIR for *ectC* |
| C(XX)Rv | TCAGACGGTTTCGGCCTCCAGC | Reverse primer for the amplification of *ectC* from *P. stutzeri* |
| D(AA)Fw | GTCTATCCGCTGGAGGCCGAAACCGTCTGAGAAAGGAGGCCCTTCAGATGCAAGCCGACCTGTATCCCTC | Forward primer 1 for the amplification of *ectD* from *P. stutzeri*. High pTIR for *ectD* |
| D(AG)Fw | GTCTATCCGCTGGAGGCCGAAACCGTCTGAGAAAGGAGGCCCTTCAGGTGCAAGCCGACCTGTATCCCTC | Forward primer 2 for the amplification of *ectD* from *P. stutzeri*. Medium pTIR for *ectD* |
| D(TG)Fw | GTCTATCCGCTGGAGGCCGAAACCGTCTGAGAATGGAGGCCCTTCAGGTGCAAGCCGACCTGTATCCCTC | Forward primer 3 for the amplification of *ectD* from *P. stutzeri*. Low pTIR for *ectD* |
| D(XX)Rv | TGCATGCCTGCAGGTCGACTCTAGAGGATCTCAGAGATACTGTTGCGGCCG | Reverse primer for the amplification of *ectD* from *P. stutzeri* |

Black letters: Gibson assembly-directed overlapping sequences; red letters: RBS sequences; blue letters: primer annealing sequences; underlined letters: nucleotides that were varied to achieve different pTIRs.

**Table S2**: pTIR values for each TU in the selected fast- and slow-growing transformants.

| Transformant | pTIR per TU (a.u.) | | | |
|---|---|---|---|---|
| | *ask* | *ectAB* | *ectC* | *ectD* |
| Control | 107 | 15373 | 961 | 1237 |
| 46 | 34052 | 355 | 116 | 2091 |
| 82 | 34052 | 355 | 116 | 2091 |
| 86 | 34052 | 355 | 239 | 2091 |
| 112 | 34052 | 355 | 239 | 2091 |
| 128 | 34052 | 355 | 116 | 2091 |
| 131 | 34052 | 355 | 239 | 2091 |

| | | | | |
|---|---|---|---|---|
| 138 | 34052 | 355 | 239 | 2091 |
| 7 | 10016 | 18324 | 2619 | 229 |
| 61 | 10016 | 18324 | 2619 | 229 |
| 101 | 10016 | 18324 | 2619 | 229 |
| 149 | 10016 | 18324 | 2619 | 229 |

Grey-shaded cells: control values; light green-shaded cells: TU-specific lowest pTIR; medium green-shaded cells: TU-specific intermediate pTIR; dark green-shaded cells: TU-specific highest pTIR.
